# Gpr37 modulates the severity of inflammation-induced GI dysmotility by regulating enteric reactive gliosis

**DOI:** 10.1101/2024.04.09.588619

**Authors:** Keiramarie Robertson, Oliver Hahn, Beatriz G. Robinson, Arwa T. Faruk, Mathangi Janakiraman, Hong Namkoong, Kwangkon Kim, Jiayu Ye, Estelle Spear Bishop, Randy A. Hall, Tony Wyss-Coray, Laren S. Becker, Julia A. Kaltschmidt

**Author notes:** Corresponding author: Laren S. Becker, Department of Medicine, Division of Gastroenterology and Hepatology, Stanford School of Medicine, Stanford, CA 94305 USA. Corresponding author: Julia A. Kaltschmidt, Department of Neurosurgery, Stanford University School of Medicine, Stanford, CA 94305 USA.

## Abstract

The enteric nervous system (ENS) is contained within two layers of the gut wall and is made up of neurons, immune cells, and enteric glia cells (EGCs) that regulate gastrointestinal (GI) function. EGCs in both inflammatory bowel disease (IBD) and irritable bowel syndrome (IBS) change in response to inflammation, referred to as reactive gliosis. Whether EGCs restricted to a specific layer or region within the GI tract alone can influence intestinal immune response is unknown. Using bulk RNA-sequencing and *in situ* hybridization, we identify G-protein coupled receptor *Gpr37*, as a gene expressed only in EGCs of the myenteric plexus, one of the two layers of the ENS. We show that Gpr37 contributes to key components of LPS-induced reactive gliosis including activation of NF-kB and IFN-y signaling and response genes, lymphocyte recruitment, and inflammation-induced GI dysmotility. Targeting Gpr37 in EGCs presents a potential avenue for modifying inflammatory processes in the ENS.

## Introduction

Enteric glial cells (EGCs) are key players in regulating gastrointestinal (GI) homeostasis; dysregulation of EGC signaling can lead to aberrant inflammatory responses, neuronal cell death, and GI dysmotility.^1^ EGC contribution to inflammation has been explored in animal models of inflammatory bowel disease (IBD), and studies have shown that EGCs become reactive in response to inflammation, a process referred to as reactive gliosis.^2,3^ Reactive gliosis was first described in astrocytes and involves direct response to inflammatory mediators, activation of pro-inflammatory pathways and secretion of both pro- and anti-inflammatory chemokines and cytokines.^4^ Reactive gliosis in astrocytes is a heterogeneous response that depends on many factors including astrocyte location within the brain.^5^ Much like the brain, the enteric nervous system (ENS) can be subdivided into functionally specific regions based on location along the GI tract (i.e., stomach, small intestine (SI), and colon), and ENS layer (myenteric plexus (MP) and submucosal plexus (SMP)).^6,7^ Though astrocytes are diverse and vary across brain regions, the extent to which region- or layer-specific EGCs in the ENS contribute to GI health and disease remains undetermined.

The ENS is essential for GI functions such as secretion, absorption, immune regulation, and motility.^6^ Regionally, the SI is responsible for absorption of nutrients from ingested food, while the colon absorbs water and salts from remaining undigested foods and drives the propulsion of waste products for elimination.^7^ Within each region, the MP and SMP control GI motility and regulate secretory motor function, respectively.^6^ Transcriptional heterogeneity of EGCs has been identified in both the SI and colon at the single-cell level^8–10^; however, methods to transcriptionally define and compare regional EGC profiles vary. Studies use the entire gut without separating the MP and SMP layers, or focus entirely on the MP^9,10^, leaving layer-specific heterogeneity unexplored.

Our knowledge of EGCs has dramatically improved in the last few decades, and evidence of enteric reactive gliosis in IBDs suggests EGCs as attractive therapeutic targets for treating GI disorders.^1,11,12^ EGC reactivity is heterogeneous, as molecular responses and disruptions of GI functions vary in a time, region, and insult-dependent manner.^12–15^ While our understanding of EGC reactivity and response to inflammation has increased substantially, it remains unknown whether region- or layer-specific EGC genes alone are sufficient to influence intestinal immune responses.

In this study, we investigate EGC transcriptional heterogeneity in both MP and SMP layers of the SI and colon. We identify *Gpr37* as an MP-specific gene expressed by a subpopulation of intraganglionic EGCs using bulk RNA-sequencing and *in situ* hybridization. In the brain, Gpr37 is thought to be involved in regulating reactive gliois; therefore, we used lipopolysaccharide (LPS) as a model of inflammation to investigate whether Gpr37 could similarly be involved in GI inflammation. We show that signaling through Gpr37 regulates fundamental signatures of reactive gliosis in the ENS. In the absence of Gpr37, many LPS-induced transcriptional programs remain dormant or are weakened, LPS-mediated lymphocyte infiltration is reduced, and LPS-induced delay in GI transit is attenuated. Our data suggest that Gpr37 contributes to inflammation-induced GI dysmotility by regulating reactive gliosis.

## Materials & Methods

### Ethical Statement

This study conformed to the National Institutes of Health Guidelines for the Care and

Use of Laboratory Animals and was approved by the Stanford University Administrative Panel on Laboratory Animal Care.

### Animals

C57BL/6,B6;CBA-Tg(Plp1^eGFP^)10Wmac/J (Strain #033357, hereafter *Plp*^*eGFP*^) were obtained from Jackson Laboratories (Bar Harbor, ME). C57BL/6,B6.129P2-*Gpr37*^*tm1Dgen*^/J (Strain #005806, hereafter *Gpr37*^−/−^) were obtained from Randy Hall (Emory University School of Medicine). All studies used both male and female mice aged 8-14 weeks. Mice were maintained on a 12:12 LD cycle and fed a standard rodent diet, containing 18% protein and 6% fat (Envigo Teklad). Food and water were provided ad libitum and mice were group housed with a maximum of five adults per cage.

### Lipopolysaccharide induced inflammation

Lipopolysaccharide (LPS) from *E. coli* O111:B4 (Cat #tlrl-eblps) was purchased from Invivogen (San Diego, CA) and was reconstituted to 5mg/ml by homogenizing in 1ml of endotoxin-free water. Mice were randomly divided into a control group (phosphate-buffered (PBS)), and treatment group (LPS), and received either a single intraperitoneal (IP) injection of LPS (2mg/kg body weight) in PBS, or PBS alone. Mice were sacrificed 2 or 24 hours following PBS or LPS injection.

### Histology

#### Tissue dissection and processing

Mice were euthanized using CO_2_ followed by cervical dislocation. The SI and colon were removed from the abdomen and placed in cold PBS on ice. Fecal matter was flushed out using cold 1X PBS and the SI was cut into 3 sections: the duodenum, jejunum, and ileum. Each section was laid flat on filter paper and cut longitudinally along the mesenteric border. Filter paper containing open intestinal segments were stacked together and placed in 4% PFA for 1.5 hours or overnight, followed by 3x 1X PBS washes for immunohistochemistry (IHC) and *in situ* hybridization, respectively. Tissue was then pinned mucosa side down on a sylgard plate and the muscularis (longitudinal and circular muscle with myenteric plexus) was peeled away. Both the mucosal (containing the submucosal plexus) and muscularis tissue were stored in PBS with 0.1% sodium azide at 4°C for up to 3 weeks for IHC or one week for *in situ* hybridization.

#### Immunohistochemistry

Segments of the SI (jejunum and ileum) >1cm in length, and colon (mid and distal) >0.5cm in length were used for IHC experiments. Staining was performed as previously described^16^, with minor modifications. For all antibody labeling, PBT contained 0.2% Triton X-100. Primary antibodies used included human anti-HuC/D (ANNA1) (1:100,000; gift from V. Lennon), rabbit anti-HuC/D (ANNA1) (1:2000; Abcam, AB9361), goat anti-Sox10 (1:1000; R&D Systems, AF2864), rabbit anti-cFos (1:1000, Synaptic Systems, 226 008) and fluorophore-conjugated secondary antibodies (Jackson Labs and Molecular Probes).

### In situ *RNA hybridization with protein co-detection*

Tissue was prepared as described in *Histology*. RNAscope *in situ* with protein co-detection was performed using Advanced Cell Diagnostics (ACD) RNAscope Multiplex Fluorescent Reagent Kit v2 (Cat# 323100) and ACD RNA-protein Co-detection Ancillary kit (Cat # 323180) as previously described.^8^ The following RNAscope probes were used: mm-Cd274-C3 (Cat# 420501), mm-Cxcl10-C2 (Cat# 408921), mm-Elavl4-C2 (Cat# 479581), mm-Gfap-C3 (Cat# 313211), mm-Gpr37-C1 (Cat# 319291), mm-Mbp-C3 (Cat# 451491), mm-Pde4b-C1 (Cat# 577191), mm-Plp1-C1 (Cat# 428181), and mm-Tacr3-C1 (Cat# 481671). The following antibodies were used: chicken anti-GFAP (1:5000, Abcam, AB4674), chicken anti-GFP (1:250, Abcam, AB13970), goat anti-Sox10 (1:500; R&D Systems, AF2864), human anti-HuC/D (ANNA1) (1:50,000; gift from V. Lennon) and rabbit-anti HuC/D (ANNA1) (1:1000; Abcam, AB9361).

### Image acquisition and analysis

Images were acquired on a Leica Sp8 confocal microscope using a 20x oil objective. Z stacks with 2um between each focal plane were acquired for 25-35um thick sections. Large tile images were collected for each tissue and stitched together using Navigation mode in the LASX software. Three random 500um x 500um sections per mouse were chosen from the large tile image for analysis. For colocalization analysis, cells were manually counted using the FIJI multiple point tool. The corrected total cell fluorescence (CTFC) and mean fluorescence intensity (MFI) were calculated as previously described. ^17,18^

### Tissue dissociation and cell isolation

Tissue was prepared as previously described.^19^ Briefly, after dissection, a glass rod was inserted through the lumen of the intestinal segment and a cotton-tipped swab moistened with cold PBS was used to peel away the muscularis under a dissecting scope. Peeled muscularis and lamina propria sections were placed on ice in cold FACs buffer until all tissue was collected. Lamina propria segments were incubated with stirring in Hank’s Buffered Saline Solution (HBSS) (GIBCO, Cat# 14170112) with 2% bovine calf serum (BCS) (Gemini, Cat# 100-504) and 2mM EDTA (Sigma-Aldrich, Cat#E8008) for 20 minutes at 37°C. Tissue was then transferred to 15ml conical tubes containing dispase (250ug/ml; STEMCELL Technologies, Cat# 07913) and collagenase XI (1mg/ml; Sigma-Aldrich, Cat# C7657), and digested for 60 minutes at 37°C. Following digestion, cell preparation and cell surface marker staining was carried out as previously described.^20^ The following antibodies were used for staining: mouse anti-CD11b (Biolegend, Cat# 101257), mouse anti-CD19 (Biolegend, Cat# 115520), mouse anti-CD3 (Biolegend, Cat# 100220), mouse anti-CD45 (Biolegend, Cat# 103132), and mouse anti-F4/80 (Biolegend, Cat# 123110).

### Cell sorting and flow cytometry

Enteric glia were sorted into QIAzol (Qiagen, from RNeasy Mini Kit, cat# 217004) (for quantitative real-time PCR [rt-qPCR]) or RNeasy Plus lysis buffer (Qiagen, from RNeasy Plus Micro Kit, Cat# 74034) (for sequencing) using BD FACSAria II. FACS analysis was carried out using BD LSRII. Cell sorting/flow cytometry analysis for this project was done on instruments in the Stanford Shared FACS Facility. Further analysis was performed using the FlowJo Software (Tree Star Inc.).

### RNA isolation

RNA was isolated for sequencing using RNeasy Plus Micro Kit (Cat# 74034).

### Bulk RNA-sequencing

cDNA and library syntheses were performed in house using the Smart-seq2 protocol as previously described^21,22^, with the following modifications: 8ul of extracted RNA was reverse-transcribed and the resulting cDNA amplified using 17 cycles. After bead cleanup using 0.7x ratio with AMPure beads (Thermo Fisher, A63881), cDNA concentration was measured using the Quant-iT dsDNA HS kit (Thermo Fisher, Q33120) and normalized to 0.4ng/ul as input for library prep. 0.8ul of each normalized sample was mixed with 1.2ul of tagmentation mix containing Tn5 Tagmentation enzyme (20034198, Illumina) and then incubated at 55°C for 12 minutes. The reaction was stopped by burying the plate in ice for 2 minutes followed by quenching with 0.8ul 0.1% sodium dodecyl sulfate (Teknova, S0180). 1.6ul indexing primer (IDT) was added and amplified using 12 cycles. Libraries were pooled and purified using two purification rounds with a ratio of 0.8x and 0.7x AMPure beads. Library quantity and quality was assessed using a Bioanalyzer (Agilent) and Qubit dsDNA HS kit (Invitrogen, Q33231). Sequencing of the resulting libraries was performed in house on an Illumina NextSeq 550 (Illumina) using single-end high-output 75bp kit (Ilumina, Cat# 20024906), aiming for 10 million 75bp reads per sample. Pipetting steps were performed using the liquid-handling robots Dragonfly or Mosquito HV (SPT Labtech) using 384 well-plates, and PCR reactions were carried out on a 384-plate Thermal Cycler (BioRad).

### RNA-sequencing analysis

Raw sequencing files were demultiplexed and known adapter sequences were trimmed with bcl2fastq. Data analysis of raw sequencing data was performed as previously described.^22^ Read quality control and counting were performed as previously described. ^23^ Raw sequence reads were trimmed to remove adaptor contamination and poor-quality reads using Trim Galore! (v.0.4.4, parameters: --paired --length 20 --phred33 --q 30). Trimmed sequences were aligned using STAR (v.2.5.3, default parameters). Multi-mapped reads were filtered. Read quality control and counting were performed using SeqMonk v.1.48.0 and RStudio v.3.6. Data visualization and analysis were performed using custom Rstudio scripts and the following Bioconductor packages: Deseq2^24^, topGO, destiny and org.Mm.eg.db. Finally, we excluded pseudogenes and predicted genes from the count matrix to focus predominantly on well-annotated, protein-coding genes. In total, all the following analyses were performed on the same set of 21,076 genes.

### Functional behavior

#### Whole GI transit time

Whole GI transit time was performed as previously described.^25^ Briefly, mice were gavaged with a 0.5% methylcellulose and 6% carmine red mixture and time was recorded until a red pellet was expelled.

#### Gastric emptying and SI transit

Gastric emptying and SI transit were determined as previously described.^25,26^ Briefly, mice were fasted overnight, and water was removed 3 hours prior to the start of the experiment. Mice were gavaged with a 2% methlycellulose and 2.5mg/mL rhodamine B dextran mixture (Invitrogen, D1841, MW: 70,000). After 15 minutes, mice were euthanized with CO_2_ and the stomach and SI were removed. The SI was then divided into 10 equals segments and all 10 segments, and the stomach were placed into a 5ml Eppendorf tube containing saline. The stomach and segments were homogenized, and fluorescence was measured using a plate reader. Gastric emptying and geometric center were calculated as previously described.^26^

#### Fecal water content

Fecal water content was determined as previously described.^27^ Percentage of water content was calculated as previously described.^25^

### Ex vivo *colonic motility monitor artificial pellet assay*

Colonic motility was assessed *ex vivo* using an adapted setup previously described.^25,28,29^ The artificial pellet assay was adapted from.^30^ Colons were removed without the cecum, and contents were carefully flushed out with warm Krebs solution. Mesentery was cut away and colons were pinned down by the proximal and distal ends in an organ bath containing warm circulating Krebs solution (NaCl, 120.9mM; KCl, 5.9mM; NaHCO_3_, 25.0mM; Monobasic NaH_2_PO_4_, 1.2mM; CaCl_2_, 3.3mM; MgCl_2_•6H20, 1.2mM; D-Glucose, 11.1mM) saturated with carbogen (95% O_2_ and 5% CO_2_). Colons were acclimated to the organ bath for 10 minutes. A lubricated 2mm 3D artificial pellet was inserted gently with a blunt ended gavage needle into the proximal-mid colon junction. Once the pellet was completely expelled, it was immediately placed back into the colon for a second trial. Colons were recorded for a minimum of 3 successful trials. Upon trial completion, empty colons were recorded for another 10 minutes. To determine pellet expulsion time and velocity, the pellet’s path was traced using FIJI plugin TrackMate (v7.6.1).

### Statistical analysis

Statistical analysis for IHC, *in situ* hybridization, and functional experiments was performed using GraphPad Prism (Version 9.4.1). Data are expressed as mean ±□SEM. Data with two groups were analyzed using unpaired two-tailed t-test. When more than 2 groups were compared, a two-way ANOVA with Tukey’s multiple comparison test was used. *P* less than 0.05 was considered significant.

## Results

### EGCs display region- and layer-specific heterogeneity

Neuronal density and organization in the adult ENS differ by intestinal regions and layers.^16^ We asked, similarly, whether EGCs also differ in their region- and layer-specific density and organization. Hamnett et al^16^ reported a higher density of glia in the colon MP as compared to the SI MP, and a higher density of glia in the MP compared to the SMP overall.^16^ To assess whether EGCs additionally differ in their region- and layer-specific organization, we used mice in which GFP was expressed under the proteolipid protein 1 (Plp1) promoter (*Plp1*^*eGFP*^), as Plp1 is the most widely expressed marker of EGCs in both the MP and SMP.^31^ We combined endogenous GFP expression to visualize EGCs with immunohistochemistry (IHC) against neuronal marker HuC/D in wholemount sections of all regions and layers of the ENS (Figure 1A). We found that EGCs mirror the region- and layer-specific heterogeneity of enteric neurons (Figure 1B).

**Figure 1:**
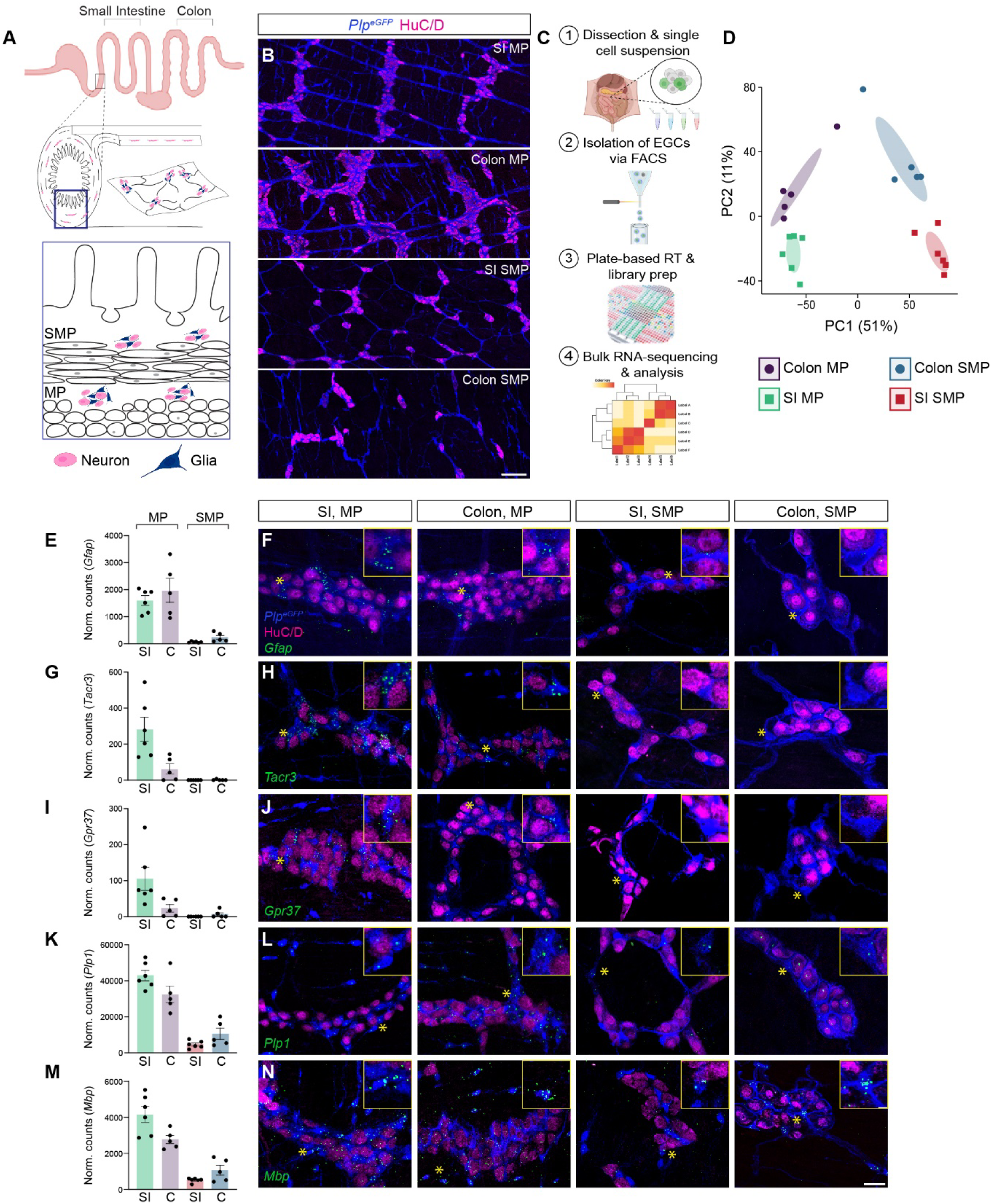
EGCs display region- and layer-specific heterogeneity. **(A)** chematic of the gastrointestinal tract regions (small intestine (SI) and colon) (top) and layers (myenteric plexus (MP), submucosal plexus (SMP) (middle) of the intestine), and a cross section of GI tract with the location of neurons (pink) and glia (blue) in the MP and SMP (bottom). **(B)** Endogenous GFP labeling of Plp1 expressing EGCs (blue) and neuronal label HuC/D (pink) in the SI and colon MP and SMP of *Plp1*^*eGFP*^ mice. Scale bar represents 100um. **(C)** Schematic of bulk-RNA sequencing experimental pipeline. **(D)** PCA of sorted EGCs from the SI MP (green squares), colon MP (purple circles), SI SMP (red squares), and colon SMP (blue circles) of *Plp1*^*eGFP*^ mice. **(E-N)** Normalized counts and corresponding representative images of the SI MP, colon MP, SI SMP and colon SMP of Plp*1*^*eGFP*^ mice for GFP to label EGCs (blue), neuronal marker HuC/D (pink) and example validation genes (green): *Gfap* (E and F), *Tacr3* (G and H), *Gpr37* (I and J), *Plp1* (K and L) and *Mbp* (M and N). Asterix represents the enlarged image in the top right corner of each image. Scale bar represents 25um, 5um for enlarged images. All charts (mean ± SEM) (n ≥5).

We next asked if EGC transcriptional profiles possess region- and/or layer-specific heterogeneity in the SI and colon. We used adult *Plp1*^*eGFP*^ mice to isolate EGCs from the MP and SMP of the SI and colon for bulk RNA-sequencing (Figure 1C). Using principal component analysis (PCA), we found that 50% of the variance is driven by the layer in which the EGCs reside (MP versus SMP). In comparison, 11% of the variance is driven by the region they reside in (SI versus colon) (Figure 1D).

We next performed *in situ* hybridization in combination with IHC on wholemount tissue of *Plp1*^*eGFP*^ mice to confirm expression of genes that we identified from the PCA to be differentially expressed in the MP versus the SMP and the SI versus the colon. Glial fibrillary acid protein *Gfap*, tachykinin receptor *Tacr3*, and G-coupled protein receptor *Gpr37*, are largely restricted to the MP (Figure 1E-J). In addition to layer differences, *Tacr3* and *Gpr37* also showed regional differences, with higher expression in the SI versus the colon. In comparison, *Plp1* and the brain oligodendrocyte gene myelin basic protein *Mbp*, were expressed in both the MP and SMP of the SI and colon (Figure 1K-N). Of note, due to immune cell contamination in our SMP data, we were unable to identify SMP-specific EGC genes with *in situ* hybridization/IHC. Taken together, our results show that EGCs in different regions and layers of the ENS are transcriptionally distinct.

### Gpr37 promotes Gfap increases in EGCs during inflammation

We decided to focus on Gpr37 as an MP-restricted EGC gene that in the CNS, has been suggested to play a protective role by regulating astrocyte reactive gliosis.^32^ EGCs can be classified into four subtypes based on morphology and location.^33^ One subtype, intraganglionic EGCs localized within enteric ganglia, have been proposed to be involved in the regulation of neuroinflammation.^1,34^ Published single-cell sequencing screens have shown Gpr37 to be expressed by a subpopulation of EGCs within the MP.^8–10^ We performed *in situ* hybridization/IHC on *Plp1*^*eGFP*^ mice in the MP and SMP of the SI and colon and found *Gpr37* selectively expressed in intraganglionic EGCs (Figure 2A).

**Figure 2:**
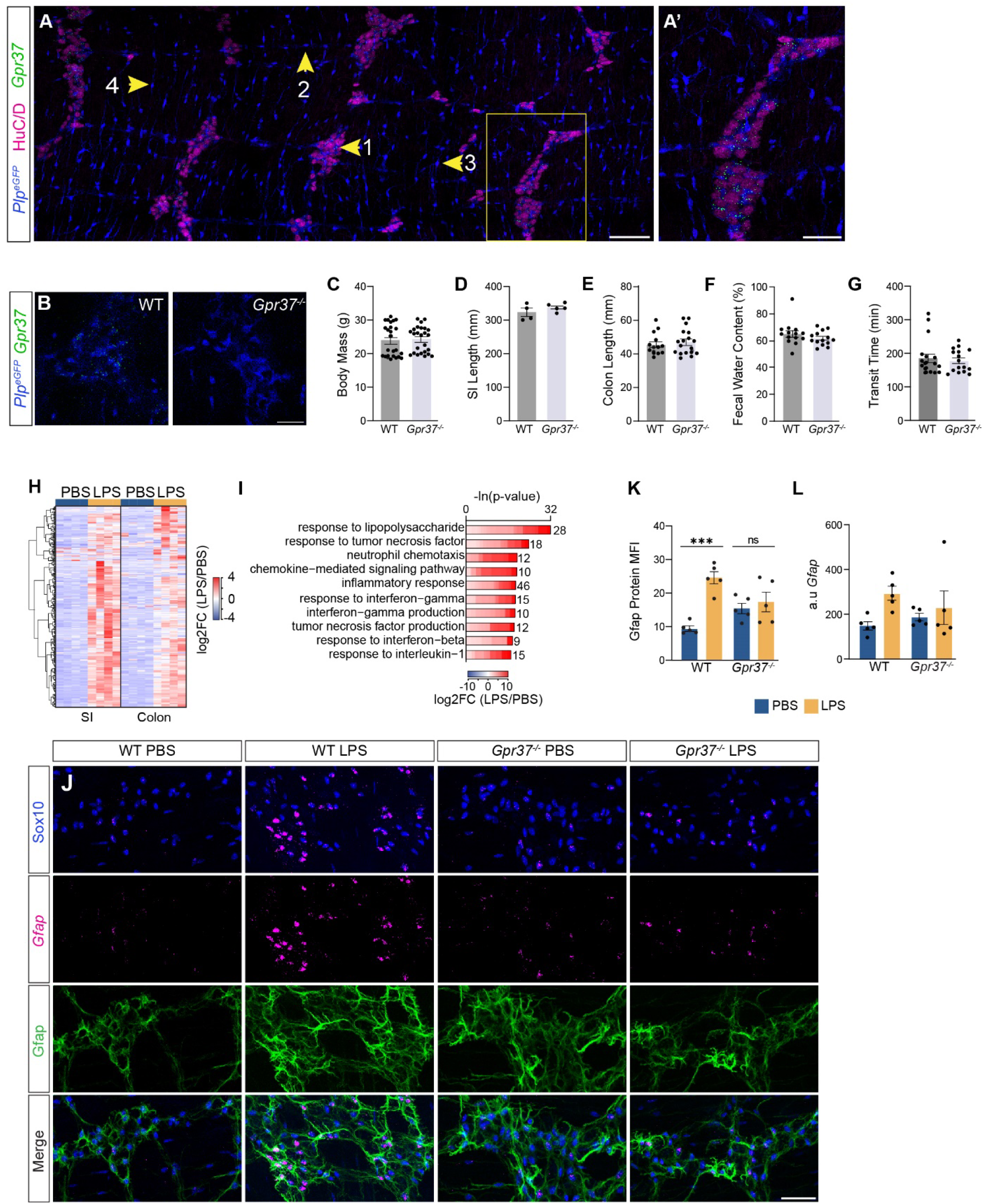
Gpr37 promotes Gfap increases in EGCs during inflammation. **(A)** Representative images of the SI MP of *Plp1*^*eGFP*^ mice for GFP to label EGCs (blue), neuronal marker HuC/D (pink), and *Gpr37* (green). Numbers correspond to EGC subtypes: (1) Intraganglionic, (2) Interganglionic, (3) Extraganglionic and (4) Intramuscular. Scale bar represents 100um. Yellow box indicates enlarged ganglia **(A’)**. Scale bar represents 50um. **(B)** Representative images of the SI MP of *Gpr37*^*WT*^; *Plp*^*eGFP*^ and *Gpr37*^−/−^; *Plp*^*eGFP*^ mice for GFP to label EGCs (blue) and *Gpr37* (green). Scale bar represents 30um. **(C-G)** Analysis of body mass (C), SI length (D), colon length (E), fecal water content (F), and whole GI transit time (G) in *Gpr37*^*WT*^ and *Gpr37*^−/−^ mice. **(H)** Heatmap of EGC genes significantly upregulated by LPS in the SI and colon MP of *Plp1*^*eGFP*^ mice (p.adj <0.05). Colors indicate gene-wise log2 fold changes. **(I)** Representative gene ontology (GO) analysis of EGC genes significantly upregulated by LPS in the SI MP. Lengths of bars represent negative ln-transformed padj using Fisher’s exact test. Colors indicate gene-wise log2 fold changes in the SI MP. **(J)** Representative images of the colon MP of *Gpr37*^*WT*^ PBS, *Gpr37*^*WT*^ LPS, *Gpr37*^−/−^ PBS and *Gpr37*^−/−^ LPS mice for Sox10 (blue) and *Gfap* (pink) and Gfap (green). Scale bar represents 40um. **(K)** Mean fluorescent intensity (MFI) analysis of Gfap protein in the colon MP of *Gpr37*^*WT*^ PBS, Gpr*37*^*WT*^ LPS, *Gpr37*^−/−^ PBS and *Gpr37*^−/−^ LPS mice. **(L)** Corrected total cellular fluorescence (CTFC) analysis of *Gfap* RNA in the colon MP of *Gpr37*^*WT*^ PBS, Gpr*37*^*WT*^ LPS, *Gpr37*^−/−^ PBS and *Gpr37*^−/−^ LPS mice. All data (mean ± SEM). (C-G) Unpaired t-test. (K-L) Two-way Anova with Tukey’s multiple comparisons test. * p<0.05; ** p<0.01; *** p<0.001; **** p<0.0001; ns, no significant difference. n ≥4 for all experiments. Data from H and I were collected 2 hours after PBS or LPS injection. Data from J, K and L were collected 24 hours after PBS or LPS injection.

To probe the role of Gpr37 in gut motility, we used a *Gpr37*^−/−^ transgenic mouse line and confirmed the absence of *Gpr37* in EGCs (Figure 2B). *Gpr37*^−/−^ mice had normal body weight, SI and colon length as compared to *Gpr37*^*WT*^ controls (Figure 2C-E). Further, fecal water content and whole GI transit time were unaffected in *Gpr37*^−/−^ mice (Figure 2F-G).

Given the association of Gpr37 with reactive gliosis in the brain and its specificity to a neuroinflammation-related EGC subtype, we next asked if Gpr37 influences EGC reactivity in intestinal inflammation.^1,32^ We used intraperitoneal injection of LPS to induce intestinal inflammation. LPS has previously been shown to activate the immune system, causing increases in pro-inflammatory cytokines and recruitment of immune cells in all layers and regions of the mouse ENS.^35^ We asked whether inflammation-associated transcriptional programs were activated in EGCs during LPS-induced inflammation. Using bulk-RNA sequencing of PBS- or LPS-injected *Plp1*^*eGFP*^ mice, we found that LPS led to the upregulation of 341 (SI) and 225 (colon) genes (Figure 2H). We further observed differential expression of LPS-induced genes in the SI versus the colon (Figure S1A). We applied gene ontology (GO) enrichment analysis to these gene sets and found enrichment of genes associated with inflammatory pathways such as “neutrophil chemotaxis”, “chemokine-mediated signaling pathway”, and “response to interferon-beta”, in the SI and colon MP (Figure 2I, Figure S1B). Together, these results suggest that EGCs respond to inflammation via the upregulation of genes involved in inflammatory processes.

We next asked if a lack of Gpr37 could affect the response of EGCs to inflammation. Increased Gfap expression indicates reactive gliosis and has been observed in the intestines of ulcerative colitis and Crohn’s patients.^3,36^ Inflammation-induced increases in astrocyte Gfap protein peak at 24 hours^37^; therefore we performed immunohistochemistry against Gfap 24 hours post PBS or LPS injection in the SI and colon of *Gpr37*^*WT*^ and *Gpr37*^−/−^ mice (Figure 2J). We found a 2.6-fold increase in Gfap protein expression in the colon of *Gpr37*^*WT*^ mice following LPS injection, but this increase was absent in LPS-injected *Gpr37*^−/−^ mice (Figure 2K). We further corroborated these experiments by measuring the total cellular fluorescence of *Gfap* RNA; following LPS injection, *Gpr37*^*WT*^ mice showed increases in *Gfap*, whereas *Gpr37*^−/−^ mice did not (Figure 2L). Our results show that the LPS-mediated increase in Gfap, a marker of reactive gliosis, is absent or attenuated in *Gpr37*^−/−^ mice.

### Transcriptional profiling of EGC inflammatory response reveals Gpr37 participates in pathways associated with reactive gliosis

We next asked whether additional immune-related transcriptional programs are attenuated in the absence of Gpr37. To fluorescently label EGCs in the absence of Gpr37, we intercrossed *Gpr37*^−/−^ with *Plp1*^*eGFP*^ mice. We isolated GFP+ EGCs from SI and colon MP of *Gpr37*^−/−^; *Plp1*^*eGFP*^ and *Gpr37*^*WT*^; *Plp1*^*eGFP*^ mice 24 hours post PBS or LPS injection and performed bulk RNA-sequencing. Differential gene expression (DEG) analysis revealed striking differences in the number of LPS-regulated genes in *Gpr37*^*WT*^; *Plp1*^*eGFP*^ versus *Gpr37*^−/−^; *Plp1*^*eGFP*^ mice. In the SI, absence of Gpr37 led to a 46% and 82% decrease of LPS-induced up- and downregulated DEGs, respectively (Figure S2A-B). In the colon, the attenuation was even more striking, with a 75% and 83% reduction in the number of LPS-induced up- and downregulated DEGs, respectively (Figure 3A-B). GO enrichment analysis revealed genes associated with cytokine signaling and production, apoptotic signaling and T cell regulation were reduced in *Gpr37*^−/−^; *Plp1*^*eGFP*^ EGCs compared to *Gpr37*^*WT*^; *Plp1*^*eGFP*^ EGCs in both the SI and colon (Figure S2C, Figure 3C). Loss of Gpr37 attenuated LPS induction of interferon signaling and response genes (Figure S2D-E), which is notable given the critical role interferon signaling in EGCs has in intestinal homeostasis and reactive gliosis.^38^ Gpr37 also appears to target the NF-kappaB (NF-kB) signaling pathway, involved in the regulation of pro-inflammatory responses^39,40^; many NF-kB target genes induced by LPS in the SI and colon in *Gpr37*^*WT*^; *Plp1*^*eGFP*^ EGCs were attenuated in EGCs of *Gpr37*^−/−^; *Plp1*^*eGFP*^ mice (Figure 3D-E). Of the NF-kB target genes found in our data set, *Cd274* (Figure 3F), *Ccl2, Cxcl10* (Figure 3G), *Gbp2*, and *Cd44*, have shown increased expression in EGCs of mice infected with *Heligmosomoides polygyrus*, a model system that has been used to study reactive gliosis in EGCs, and are upregulated in patients with ulcerative colitis.^41^ We validated our findings with *in situ* hybridization using co-labeling with Sox10 as a marker for EGCs^31^, and found that *Cxcl10* and *Cd274* are upregulated by LPS in colonic EGCs of *Gpr37*^*WT*^; *Plp1*^*eGFP*^ mice, but not in the absence of Gpr37 (Figure 3I-L).

**Figure 3:**
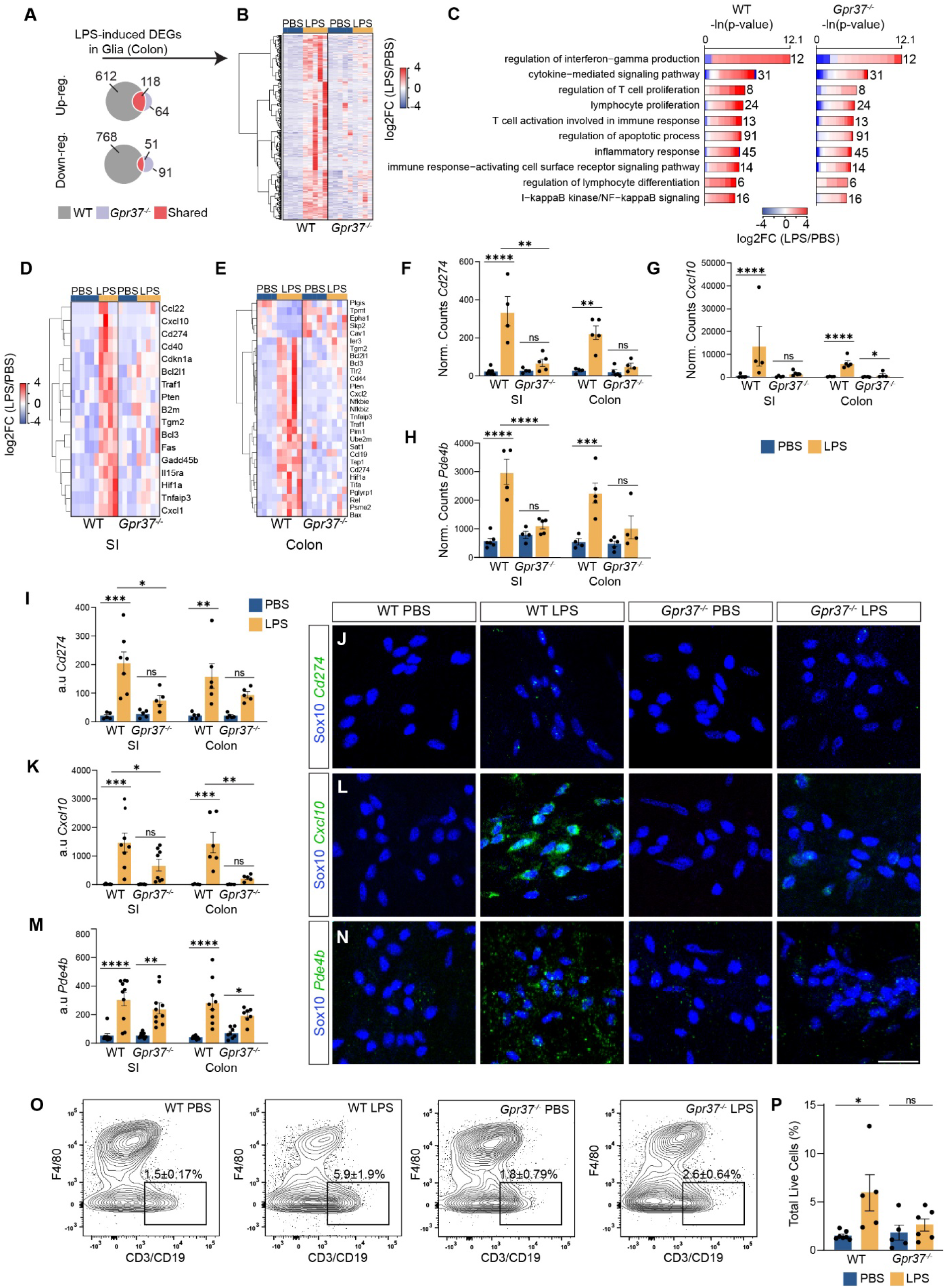
Transcriptional profiling of EGC inflammatory response reveals Gpr37 participates in pathways associated with reactive gliosis. **(A)** Venn diagram of LPS-induced differentially expressed genes (DEGs) in colonic EGCs. Gray indicates DEGs in Gpr*37*^*WT*^; *Plp1*^*eGFP*^ mice only, purple indicates DEGs in *Gpr37*^−/−^; *Plp1*^*eGFP*^ mice only, and red indicates shared DEGs between Gpr*37*^*WT*^; *Plp1*^*eGFP*^ *and Gpr37*^−/−^; *Plp1*^*eGFP*^ mice. **(B)** Heatmap of all LPS-induced upregulated DEGs in the colon MP. Colors indicate gene-wise log2 fold changes. **(C)** Inflammatory related GO terms of the 612 LPS-induced upregulated DEGs in Gpr*37*^*WT*^; *Plp1*^*eGFP*^ colonic MP EGCs only. Lengths of bars represent negative ln-transformed padj using Fisher’s exact test. Colors indicate gene-wise log2 fold changes in the colon MP. **(D and E)** Heatmap of differentially expressed NF-kB target genes in the SI MP (D) and the colon MP (E). Colors indicate gene-wise log2 fold changes. **(F-H)** *Cd274* (F), *Cxcl10* (G), and *Pde4b* (H) normalized counts in the SI and colon MP of *Gpr37*^*WT*^ PBS, Gpr*37*^*WT*^ LPS, *Gpr37*^−/−^ PBS, and *Gpr37*^−/−^ LPS mice. **(I, K, M)** Corrected total cellular fluorescence (CTFC) of *Cd274* (I), *Cxcl10* (K), and *Pde4b* (M), in the SI and colon MP of *Gpr37*^*WT*^ PBS, Gpr*37*^*WT*^ LPS, *Gpr37*^−/−^ PBS, and *Gpr37*^−/−^ LPS mice. **(J, L, N)** Representative images of *Cd274* in the SI MP (J), *Cxcl10* in the colon MP (L), and *Pde4b* in the colon MP (N), of *Gpr37*^*WT*^ PBS, *Gpr37*^*WT*^ LPS, *Gpr37*^−/−^ PBS, and *Gpr37*^−/−^ LPS mice. Images are labeled with Sox10 (blue) and *Cd274, Cxcl10, and Pde4b* (green). Scale bar represents 25um. **(O)** Representative flow cytometric plots of single CD45+ live cells from the colon MP of *Gpr37*^*WT*^ PBS, *Gpr37*^*WT*^ LPS, *Gpr37*^−/−^ PBS and *Gpr37*^−/−^ LPS mice using cell surface markers F4/80, CD3 and CD19. Boxes represent lymphocytes (CD3/CD19+). Numbers on plot represent percent (mean±SEM) of live cells. **(P)** Proportion of colonic MP lymphocytes (CD45+CD3/CD19+) expressed as a percent of live cells for *Gpr37*^*WT*^ PBS, Gpr*37*^*WT*^ LPS, *Gpr37*^−/−^ PBS and *Gpr37*^−/−^ LPS mice. All data (mean ± SEM). (I, K, M, P) Two-way Anova with Tukey’s multiple comparisons test. * p<0.05; ** p<0.01; *** p<0.001; **** p<0.0001; ns, no significant difference. n ≥4 for all experiments. All experiments were conducted 24 hours after LPS or PBS injection.

During inflammation, increased levels of phosphodiesterases lead to intracellular cAMP degradation, and release of NF-kB inhibition, allowing for a pro-inflammatory response.^42^ Pde4b is a phosphodiesterase expressed by many immune cells as well as EGCs.^43^ Our sequencing data revealed that in response to LPS, *Pde4b* was significantly increased in EGCs in both the SI and colon of *Gpr37*^*WT*^; *Plp1*^*eGFP*^ mice, however, this increase was lost in the absence of Gpr37 (Figure 3H). Validation with *in situ* hybridization/IHC showed an increase in *Pde4b* after LPS in EGCs of *Gpr37*^*WT*^ mice, which was attenuated in LPS-treated *Gpr37*^−/−^; *Plp1*^*eGFP*^ mice (Figure 3M-N). These results indicate that Gpr37 regulates gene expression changes associated with inflammation.

We next asked whether loss of Gpr37 affects immune cell composition following LPS treatment. Using flow cytometry, we found a 4-fold increase in lymphocyte (CD45+CD3/CD19+ cells) infiltration in the colon MP of LPS-treated *Gpr37*^*WT*^; *Plp1*^*Egfp*^ mice. In comparison, we found only a 1.5-fold increase of colonic lymphocytes in LPS-treated *Gpr37*^−/−^; *Plp1*^*eGFP*^ mice (Figure 3O-P), indicating Gpr37 deficiency attenuates LPS-induced lymphocyte recruitment. Total lymphocytes in the SI trended lower in *Gpr37*^−/−^ mice, regardless of PBS or LPS (Figure S2F). While monocytes (CD45+F4/80-CD11b+ cells) trended higher with LPS, loss of Gpr37 had no significant effect on the increased trend in either the SI or colon (Figure S2G-H). Similarly, total leukocytes (CD45+ cells) and macrophages (CD45+F4/80+CD11b+ cells) trended higher with LPS in the colon, and loss of Gpr37 did not significantly affect this trend (Figure S2I-K). Total leukocytes and macrophages were unaffected by LPS in the SI, regardless of the presence or absence of Gpr37 (Figure S2L-M).

Taken together, these results show that Gpr37 plays a role in enteric reactive gliosis, partially through influences on NF-kB and interferon signaling and response genes, and lymphocyte recruitment.

### Gpr37 contributes to changes in neuronal activity and inflammation-induced GI dysmotility

Changes in the physiological properties of neurons during inflammation have been linked to GI dysmotility.^44^ We next asked whether Gpr37 is involved in the activation of neurons during inflammation. We performed IHC against the activity-dependent marker cFos in the SI and colon MP of *Gpr37*^*WT*^ and *Gpr37*^−/−^ mice two hours after PBS or LPS injection.^45^ We found a two-fold increase in neuronal cFos expression in the distal colon of LPS-treated *Gpr37*^*WT*^ mice, as compared to a 1.5-fold increase in LPS-treated *Gpr37*^−/−^ mice, suggesting Gpr37 plays a role in regulating neuronal activity during inflammation (Figure 4A-B).

**Figure 4:**
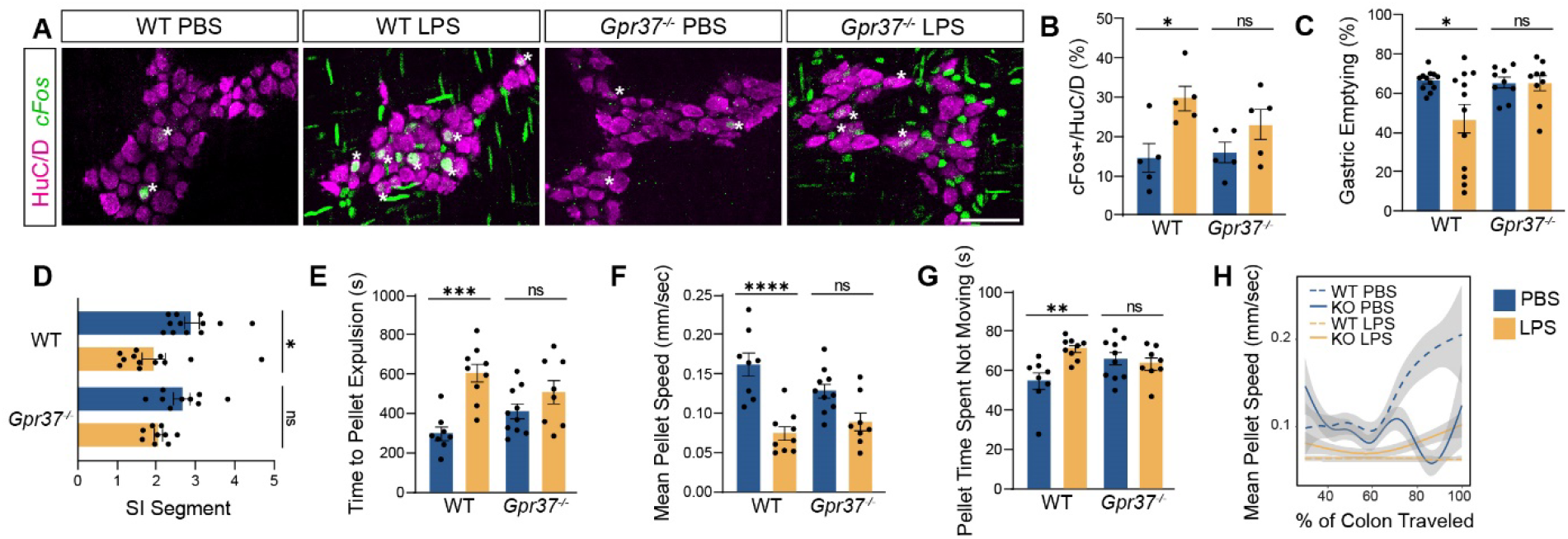
Gpr37 contributes to changes in neuronal activity and inflammation-induced GI dysmotility. **(A)** Representative images of the colon MP of *Gpr37*^*WT*^ PBS, *Gpr37*^*WT*^ LPS, *Gpr37*^−/−^ PBS, and *Gpr37*^−/−^ LPS mice for neuronal marker HuC/D (pink) and cFos (green). Asterix indicates cells co-expressing HuC/D and cFos. Scale bars represent 50um. **(B)** Proportion of total HuC/D cells expressing cFos in the distal colon MP of*Gpr37*^*WT*^ PBS, *Gpr37*^*WT*^ *LPS, Gpr37*^−/−^ PBS, and *Gpr37*^−/−^ LPS mice. **(C)** Stomach motility quantified as percent gastric emptying in *Gpr37*^*WT*^ PBS, *Gpr37*^*WT*^ LPS, *Gpr37*^−/−^ PBS, and *Gpr37*^−/−^ LPS mice. **(D)** SI motility quantified as geometric center for *Gpr37*^*WT*^ PBS, *Gpr37*^*WT*^ LPS, *Gpr37*^−/−^ PBS, and *Gpr37*^−/−^ LPS mice. **(E)** Time it takes for the artificial pellet to expel from the colon of *Gpr37*^*WT*^ PBS, *Gpr37*^*WT*^ LPS, *Gpr37*^−/−^ PBS, and *Gpr37*^−/−^ LPS mice. **(F)** Mean artificial pellet speed as it travels through the colon of *Gpr37*^*WT*^ PBS, *Gpr37*^*WT*^ LPS, *Gpr37*^−/−^ PBS, and *Gpr37*^−/−^ LPS mice. **(G)** Percentage of time the artificial pellet spends not moving in the colon of *Gpr37*^*WT*^ PBS, *Gpr37*^*WT*^ LPS, *Gpr37*^−/−^ PBS, and *Gpr37*^−/−^ LPS mice. **(H)** Local Regression (LOESS) of mean pellet speed for *Gpr37*^*WT*^ PBS, *Gpr37*^*WT*^ LPS, *Gpr37*^−/−^ PBS, and *Gpr37*^−/−^ LPS mice as a function of the % colon length traveled. 95% confidence interval shown in gray. All data (mean ± SEM). (B-G) Two-way Anova with Tukey’s multiple comparisons test. * p<0.05; ** p<0.01; *** p<0.001; **** p<0.0001; ns, no significant difference. n ≥5 for all experiments. Data from A and B was collected 2 hours after PBS or LPS injection. All other data was collected 24 hours after PBS or LPS injection.

Preventing EGC reactivity has been shown to restore inflammation-induced changes in GI function.^46–48^ Given that signatures of reactive gliosis and LPS-induced increases in neuronal cFos are absent or attenuated in the absence of Gpr37, we next asked whether loss of Gpr37 influences inflammation-induced GI dysmotility. We assessed *in vivo* stomach and SI transit in PBS- and LPS-treated *Gpr37*^*WT*^ and *Gpr37*^−/−^ mice by measuring gastric emptying and geometric center.^49^ We found that LPS decreased gastric emptying and geometric center by 30% and 33% in *Gpr37*^*WT*^ mice, but only by 1% and 23% in *Gpr37*^−/−^ mice (Figure 4C-D). These results indicate that loss of Gpr37 attenuates LPS-induced disruption of stomach and SI motility.

We next assessed *ex vivo* colonic activity in PBS- and LPS-injected *Gpr37*^*WT*^ and *Gpr37*^−/−^ mice. We inserted a 3D-printed artificial fecal pellet into the proximal/mid-colon junction and recorded the time to expulsion. We found that LPS treatment doubled the time to pellet expulsion in *Gpr37*^*WT*^ mice, while showing no significant increase in expulsion time in *Gpr37*^−/−^ mice (Figure 4E). Furthermore, LPS treatment resulted in a 54% decrease in pellet speed in *Gpr37*^*WT*^ mice, compared to a 31% decrease in *Gpr37*^−/−^ mice (Figure 4F). Artificial pellets spent significantly more time stationary and had overall fewer pellet movements in LPS-treated *Gpr37*^*WT*^ compared to *Gpr37*^−/−^ mice (Figure 4G-H). Taken together these results suggest that enteric glial Gpr37 regulates the severity of LPS-induced GI dysmotility.

## Discussion

In this study, we find that enteric glia possess region- and layer-specific transcripts, and that Gpr37, a known regulator of reactive gliosis in the brain^32^, is restricted to an intraganglionic population of EGCs in the MP of both the SI and colon. In response to LPS-induced inflammation, Gpr37 regulates components of the reactive enteric glia phenotype, including NF-kB and interferon signaling and response genes, lymphocyte recruitment, neuronal activation, and GI motility. Our findings were observed in both sexes; however, we cannot exclude the possibility that we were underpowered to detect subtle differences between males and females.

The composition of both neurons and immune cells varies across different regions of the GI tract, likely tailored to each location’s specific function.^16,50^ We additionally found EGC genes specific for the MP, as well as EGC genes that are differentially expressed in distinct intestinal regions. In recent decades, EGCs have come to be recognized as a diverse population, with their heterogeneity commonly defined based on morphology and location within a layer.^33^ Our knowledge of how these subtypes contribute to GI function is limited. However, it has been suggested that interganglionic EGCs, residing along nerve fibers in between ganglia, play a role in signal propagation and neuromodulation, while intraganglionic EGCs within ganglia and in close proximity to neurons, play a role in both neuromodulation and neuroinflammation.^1^ Furthermore, immune infiltration into the MP in IBDs has been shown to preferentially target ganglia.^51,52^ It is therefore plausible that intraganglionic EGCs might play a role in the preferential recruitment of immune cells observed in IBDs, as EGCs are known to communicate directly with immune cells.^36,53^ Our finding that Gpr37 is restricted to intraganglionic EGCs of the MP and regulates key features of LPS-induced EGC reactive gliosis, including immune cell recruitment, provides compelling evidence for the inflammatory role of intraganglionic EGCs.

Our work demonstrates that Gpr37 contributes to LPS-induced reactive gliosis. We found that previously described biological pathways of reactive gliosis^4,46^ matched with GO enrichment terms that were attenuated in the absence of Gpr37, such as NF-kB and IFN-y signaling pathways. Various components and downstream targets of these pathways in EGCs can influence immune cell activation, neuronal activity, and GI function during reactive gliosis.^41,54^ However, EGC reactivity is highly context-dependent.^1^ Previous work has shown that while both ATP and Il1B induce a reactive phenotype in cultured mouse EGCs, Il1B stimulation results in a more robust induction of several chemokines as compared to ATP stimulation, indicating that reactive gliosis in EGCs is not a binary response.^46^ Our work demonstrates that LPS administration induces a form of reactive gliosis, and that Gpr37 contributes to various components of the LPS-induced reactive phenotype. It will be interesting for future studies to explore whether Gpr37 regulates reactive gliosis equivalently for alternate stressors such as intestinal manipulation or DSS colitis.

EGC reactive gliosis is thought to cause GI dysmotility during inflammation through influences on neuronal circuitry.^47,55^ Our observation that LPS increases neuronal activity and delays GI transit, and that in the absence of Gpr37, these changes are attenuated, suggests Gpr37 signaling affect GI motility via its influence on neuronal activity. Whether Gpr37 interacts directly or indirectly with neurons is unknown. Pro-inflammatory mediators in EGCs, such as Il1b and Il6, have been shown to amplify neuronal excitability by altering signaling pathways in both EGCs and neurons, while others, including Cxcl2 and Cxcl10, two chemokines we found to be regulated by Gpr37, may indirectly influence neuronal activity through actions on surrounding immune cells.^1^ We found that LPS-induced lymphocyte recruitment was attenuated in the absence of Gpr37; suggesting that Gpr37 could regulate inflammation-induced dysmotility by indirectly altering neuronal activity via the secretion of cytokines and chemokines that specifically recruit lymphocytes. Changes in lymphocyte numbers have been observed in various GI disorders, and increased infiltration of lymphocytes into the MP has been suggested as a link between inflammation and motility dysfunction in IBS patients.^56^ Colonic inertia, a motility disorder that results in severe constipation, often refractory to laxatives^57^, has shown increased infiltration of lymphocytes specifically into the ganglia of the MP.^58^ Furthermore, specific MP inflammatory infiltrates have also been observed for ulcerative colitis and Crohn’s disease.^52^

Reactive gliosis is considered both a beneficial and harmful process depending on the type and severity of insult. Altering interferon signaling in EGCs during infections prevents EGCs from maintaining appropriate levels of immune cells essential for tissue repair, ultimately resulting in larger areas of tissue damage long after an inflammatory response has subsided.^41^ On the other hand, preventing the ability of EGCs to take on a reactive phenotype by disrupting their ability to respond to ATP protects against inflammation-induced neuronal cell death and delayed GI transit.^1,47^ Intact inflammatory responses are crucial for the resolution of inflammation and EGCs are key mediators in returning injured tissue to a homeostatic state.^41,60,61^ In the CNS, the role of Gpr37 in reactive gliosis supports the notion that reactive gliosis is a beneficial process, as ischemic damage is amplified in its absence^32,61^; however, whether the contribution of Gpr37 to reactive gliosis in the ENS is harmful or beneficial remains unknown. Our results show that Gpr37 contributes to pro-inflammatory outcomes and that in its absence, inflammation-induced GI dysmotility is attenuated. However, an interesting question remains whether inflammation-induced GI dysmotility is harmful or beneficial in the resolution of inflammation, or simply a side effect of a heightened immune response. Future work should focus on Gpr37’s contribution to reactive gliosis at various times throughout the inflammatory process. Moreover, given that several potential endogenous ligands/modulators of Gpr37 have been reported, including prosaposin^59^, neuroprotection D1^62^, and osteocalcin^63^, it may be of interest to determine whether these ligands can influence Gpr37 regulation of reactive gliosis in the GI tract.

EGCs are an attractive cell type for therapeutic interventions given their capacity to modulate both neuronal activity and immune function in the ENS.^11,36,64^ Increases in Gfap, a common hallmark of reactive gliosis, have been observed in IBD intestinal tissue.^3,36^ Many of the genes we found to be regulated by Gpr37 during LPS-induced inflammation also show increased expression in EGCs of patients with ulcerative colitis.^63^ Traditional IBD treatments have focused on targeting components or regulators of the inflammatory response, such as JAK inhibitors and anti-TNF agents.^65^ Not all immune cells utilizing these inflammatory pathways likely contribute to the overproduction of cytokines seen in inflammation, and non-specific anti-inflammatory treatments could be harmful to these intact processes. Furthermore, anti-inflammatory drugs have significant immunosuppressive properties, increasing the risk of infections and cancer.^66^ Gpr37-regulated reactive gliosis in EGCs could offer a more precise and less off-target approach to treating GI disorders.

## Resource Availability

### Lead Contact

Further information and requests for resources and reagents should be directed to and will be fulfilled by the lead contact, Julia Kaltschmidt.

### Materials Availability

This study did not generate new unique reagents.

### Data Availability

The sequencing datasets analyzed during the current study will be available in the Gene Expression Omnibus repository under accession number GSE262418 upon publication.

## Acknowledgements

We thank members of the Kaltschmidt laboratory for experimental advice and discussions, Vanda Lennon (Mayo Clinic) for HuC/D primary antibody, Julietta Gomez-Frittelli and Christine Plant for feedback on the manuscript and the Stanford Shared FACS Facility for their equipment. We thank Rhian Stavely and Richard Guyer for their experimental and analysis advice.

## Author contributions

Conceptualization: KR, LSB, JAK

Resources: RAH, TWC, LSB, JAK

Methodology: KR, OH, LSB, JAK

Investigation: KR, OH, BR, ATF, MJ, HN, KK, JY, ESB

Formal Analysis: KR, OH

Visualization: KR, OH

Writing-Original Draft: KR, OH, RAH, LSB, JAK

Funding Acquisition: LSB, JAK

## Declaration of generative AI and AI-assisted technologies in the writing process

During the preparation of this work the author’s used ChatGPT3.5 2022 for proofreading parts of the introduction and discussion section. After using this tool, we reviewed and edited the content as needed and take full responsibility for the content of the publication.

