## Supplemental Figures for "Gpr37 modulates the severity of inflammation-induced GI dysmotility by regulating enteric reactive gliosis"

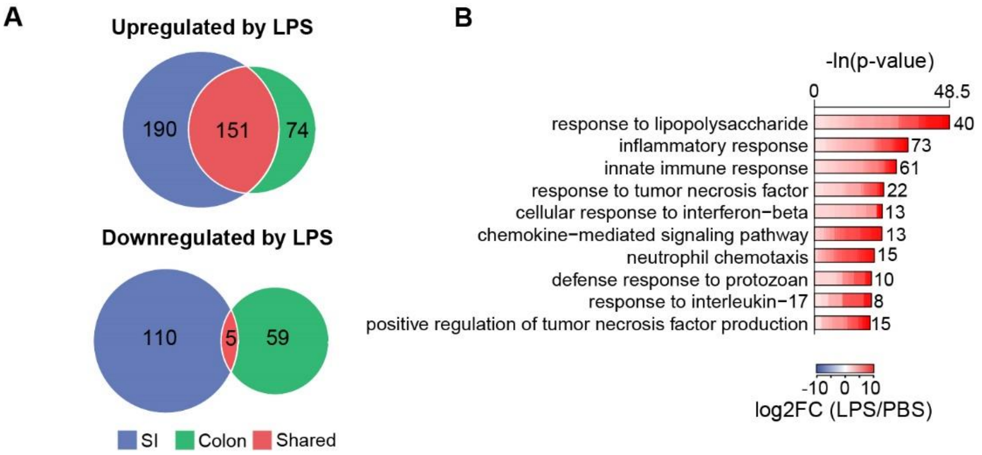


**Supplemental Figure 1: LPS induces gene expression changes in EGCs.**

**(A)** Venn diagram of LPS-induced differentially expressed genes (DEGs) in SI MP EGCs of *Plp1*$^{eGFP}$ mice. Blue indicates DEGs in the SI MP only, green indicates DEGs in the colon MP only, and red indicates shared DEGs between the SI and colon MP.

**(B)** Representative gene ontology (GO) analysis of significantly upregulated EGC genes in the colon MP of *Plp1*$^{eGFP}$ mice. Lengths of bars represent negative ln-transformed padj using Fisher's exact test. Colors indicate gene-wise log2 fold changes in the colon MP**.**

(n=4). Experiment was conducted 2 hours after PBS or LPS injection.

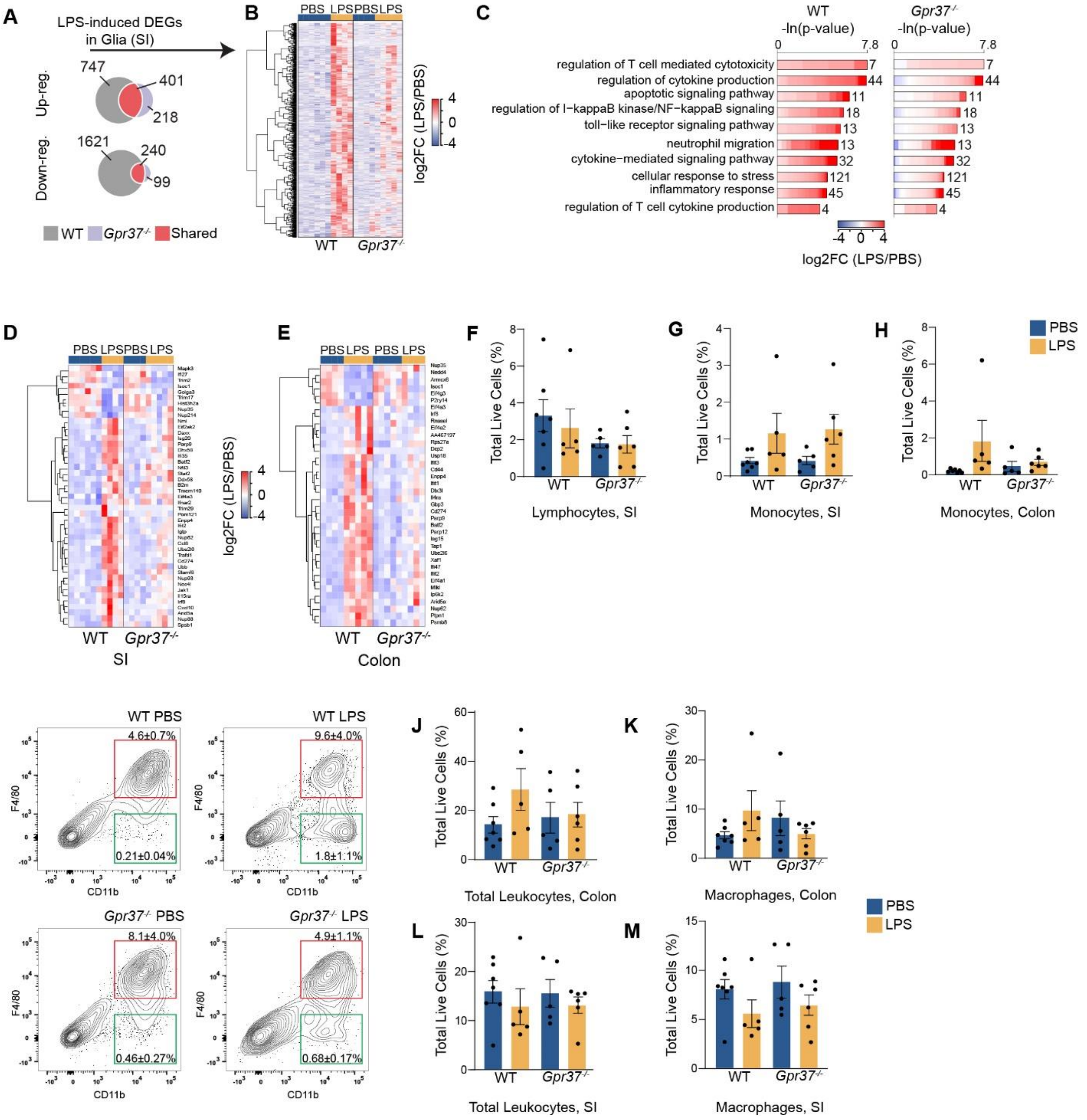


**Supplemental Figure 2: Bulk-RNA sequencing and flow cytometry analysis of LPS-induced inflammation in *Gpr37*$^{WT}$ and *Gpr37*$^{-/-}$ EGCs.**

**(A)** Venn diagram of LPS-induced DEGs in SI EGCs. Gray indicates DEGs in Gpr*37*$^{WT}$*; Plp1*$^{eGFP}$ mice only, purple indicates DEGs in *Gpr37*$^{-/-}$*; Plp1*$^{eGFP}$ mice only, and red indicates shared DEGs between Gpr*37*$^{WT}$*; Plp1*$^{eGFP}$ *and Gpr37*$^{-/-}$*; Plp1*$^{eGFP}$ mice.

**(B)** Heatmap of all LPS-induced upregulated DEGs in the SI MP. Colors indicate gene-wise log2 fold changes.

**(C)** Inflammatory related GO terms of the 747 LPS-induced upregulated DEGs in Gpr*37*$^{WT}$*; Plp1*$^{eGFP}$ SI MP EGCs only. Lengths of bars represent negative ln-transformed padj using Fisher's exact test. Colors indicate gene-wise log2 fold changes.

**(D and E)** Heatmap of differentially expressed interferon signaling genes in the SI MP (D) and the colon MP (E). Colors indicate gene-wise log2 fold changes.

**(F)** Proportion of SI MP lymphocytes (CD45+CD3/CD19+) expressed as a percent of live cells in *Gpr37*$^{WT}$ PBS, Gpr*37*$^{WT}$ LPS, *Gpr37*$^{-/-}$ PBS and *Gpr37*$^{-/-}$ LPS mice.

**(G and H)** Proportion of monocytes (CD45+F4/80-Cd11b+) in the SI MP (G) and the colon MP (H) expressed as a percent of live cells of *Gpr37*$^{WT}$ PBS, Gpr*37*$^{WT}$ LPS, *Gpr37*$^{-/-}$ PBS, and *Gpr37*$^{-/-}$ LPS mice.

**(I)** Representative flow cytometric plots of single CD45+ live cells from the colon MP of *Gpr37*$^{WT}$ PBS, *Gpr37*$^{WT}$ LPS, *Gpr37*$^{-/-}$ PBS, and *Gpr37*$^{-/-}$ LPS mice using cell surface markers F4/80 and Cd11b. Red boxes represent macrophages (F4/80+Cd11b+), and green boxes represent monocytes (F4/80-Cd11b+). Numbers on plot represent percent (mean±SEM) of live cells.

**(J-M)** Proportion of colon MP leukocytes (CD45+) (J), colon MP macrophages (CD45+F4/80+Cd11b+) (K), SI MP leukocytes (L) and SI MP macrophages (M) expressed as percent live cells in *Gpr37*$^{WT}$ PBS, Gpr*37*$^{WT}$ LPS, *Gpr37*$^{-/-}$ PBS, and *Gpr37*$^{-/-}$ LPS mice.

n ≥5. All experiments were conducted 24 hours after PBS or LPS injection.
